# Genetic deficiency of ribosomal rescue factor HBS1L causes retinal dystrophy associated with Pelota and EDF1 depletion

**DOI:** 10.1101/2023.10.18.562924

**Authors:** Shiyu Luo, Bilal Alwattar, Qifei Li, Kiran Bora, Alexandra K. Blomfield, Jasmine Lin, Anne Fulton, Jing Chen, Pankaj B. Agrawal

**Author notes:** To whom correspondence should be addressed at: Pankaj B. Agrawal, MD, Division of Neonatology, Department of Pediatrics, University of Miami Miller School of Medicine and Holtz Children’s Hospital, Jackson Health System, Miami, FL, USA.

## Abstract

Inherited retinal diseases (IRDs) encompass a genetically diverse group of conditions in which mutations in genes critical to retinal function lead to progressive loss of photoreceptor cells and subsequent visual impairment. A handful of ribosome-associated genes have been implicated in retinal disorders alongside neurological phenotypes. This study focuses on the *HBS1L* gene, encoding HBS1 Like Translational GTPase which has been recognized as a critical ribosomal rescue factor. Previously, we have reported a female child carrying biallelic *HBS1L* mutations, manifesting growth restriction, developmental delay, and hypotonia. In this study, we describe her ophthalmologic findings, compare them with the *Hbs1l*^tm1a/tm1a^ hypomorph mouse model, and evaluate the underlying microscopic and molecular perturbations. The patient was noted to have impaired visual function observed by electroretinogram (ERG), with dampened amplitudes of a- and b-waves in both rod- and cone-mediated responses. *Hbs1l*^tm1a/tm1a^ mice exhibited profound retinal thinning of the entire retina, specifically of the outer retinal photoreceptor layer, detected using in vivo imaging of optical coherence tomography (OCT) and retinal cross sections. TUNEL assay revealed retinal degeneration due to extensive photoreceptor cell apoptosis. Loss of HBS1L resulted in comprehensive proteomic alterations in mass spectrometry analysis, with169 proteins increased and 480 proteins decreased including many critical IRD-related proteins. GO biological process and GSEA analyses reveal that these downregulated proteins are primarily involved in photoreceptor cell development, cilium assembly, phototransduction, and aerobic respiration. Furthermore, apart from the diminished level of PELO, a known partner protein, HBS1L depletion was accompanied by reduction in translation machinery associated 7 homolog (Tma7), and Endothelial differentiation-related factor 1(Edf1) proteins, the latter of which coordinates cellular responses to ribosome collisions. This novel connection between HBS1L and ribosome collision sensor (EDF1) further highlights the intricate mechanisms underpinning ribosomal rescue and quality control that are essential to maintain homeostasis of key proteins of retinal health, such as rhodopsin.

## Introduction

Maintenance of cellular vitality necessitates the continuous protein synthesis by the ribosomes. However, various factors such as faulty mRNA production, inadequate availability of charged tRNAs, and genetic errors, can disrupt this process and result in detrimental cellular effects (1, 2). To mitigate these defects, organisms have evolved dedicated surveillance pathways designed to identify and prevent the accumulation of aberrant RNAs and proteins (3, 4). One such mechanism is ribosome-associated quality control (RQC) pathway, which encompasses two sequential steps, each addressing a distinct defect (5). The initial step (termed rescue) involves sensing stalled ribosomes and facilitating the splitting of 80S subunits into 40S and 60S ribosomal subunits. The subsequent step identifies 60S subunits obstructed with peptidyl-tRNA and facilitates the resolution of this aberrant structure. This leads to the release of free, translation-competent 60S ribosomal subunits and triggers the proteolysis of the nascent chain. During this process, efficient dissociation of stalled ribosomal subunits by rescue factors is crucial for optimal recruitment of downstream RQC components and nascent protein ubiquitination (6). The significance of this process in maintaining proteomic integrity and cellular fitness is underscored by the association of RQC machinery defects with various neurological diseases, such as Alzheimer’s disease and Huntington’s disease (7–10).

Rescue factor Hbs1 (Hsp70 subfamily B suppressor 1), an ortholog of mammalian HBS1L in yeast, was originally identified for its ability to salvage stalled translation in yeast strains with reduced heat shock protein 70 activity (11). Human HBS1L, a paralogue of the canonical termination factor eRF3, is expressed ubiquitously and belongs to the GTP-binding elongation factor family (12). Alternative splicing of *HBS1L* generates transcripts encoding two distinct proteins. The full length HBS1L (referred to as HBS1L or Hbs1l henceforth) interacts with pelota (PELO) to facilitate ribosomal release from stalled translation units (13). In contrast, the shorter isoform encoded by the first 4 exons of full-length HBS1L and a unique last exon (‘exon 5a’) positioned between exon 4 and exon 5 of the *HBS1L* locus, interacts with the SKI complex and functions in global mRNA turnover (14).

In previous studies, we have identified a female patient with biallelic *HBS1L* mutations leading to the loss of its longer isoforms, resulting in a phenotype characterized by severe intrauterine growth restriction, microcephaly, axial hypotonia, lax joints, global developmental delay, fused C2-C3 vertebrae, scoliosis, submucous cleft palate and retinal pigmentary deposits (15). Subsequently, we showed that *Hbs1l* hypomorph mice (*Hbs1l*^tm1a/tm1a^, with residual expression of wild-type *Hbs1l* mRNA less than 10%) recapitulates several human phenotypes, including growth restriction, facial dysmorphism, and retinal pigmentation deposit (16). Additional mouse studies show that *Hbs1l* is required for mammalian embryogenesis and complete loss of *Hbs1l* results in embryonic lethality (17). We and others have shown that the deletion of Hbs1l leads to a tissue-specific reduction in its partner protein Pelo, which alters transcription regulation and reprograms the translatome (16, 17). While both proteins are critical for cerebellar neurogenesis in mice, they are dispensable for neuronal survival post-development (17).

Inherited retinal diseases (IRDs) encompass a genetically diverse group of conditions in which mutations in genes critical for retinal function are associated with progressive demise of photoreceptor cells and associated vision loss (18). Currently, over 260 IRDs-associated genes have been identified. While most mutated genes are expressed specifically in the retina, a few ribosome-related genes (*RPL10, GTPBP1* and *GTPBP2*) have been identified along with neurological manifestations (19–21). In this study, we present the ophthalmologic findings observed in both the human patient carrying *HBS1L* mutations and *Hbs1l*^tm1a/tm1a^ hypomorph mice, demonstrating that Hbs1l deficiency triggers extensive apoptosis of photoreceptor cells as early as in 2 weeks old mice and subsequent retinal degeneration. Comprehensive proteomic analysis illustrates that the loss of Hbs1l is associated with increase in levels of 169 proteins (p-value_≤_0.05 and_≥_1.5-fold change) and reduction in 480 proteins (p-value_≤_0.05 and ≤ 0.67-fold change). GO biological process and GSEA analyses reveal that these reduced proteins are primarily enriched for: photoreceptor cell development, cilium assembly, phototransduction, and aerobic respiration. Furthermore, apart from impacting its partner protein Pelo, Hbs1l depletion is accompanied with reduced levels of translation machinery associated 7 homolog (TMA7), and Endothelial differentiation-related factor 1(EDF1) proteins. While TMA7 is closely related to cell proliferation and autophagy (22), EDF1 is known to coordinate cellular responses to ribosome collisions (23, 24), when ribosomes slow down or completely stall when they encounter obstacles on mRNAs. Together these findings suggest a broad impact by the loss of Hbsl1 and its associated transcriptional regulators on key proteins for photoreceptor health and visual function.

## Results

### Retinal dystrophy in a child carrying *HBS1L* biallelic mutations

We previously described a female child carrying compound heterozygous deleterious mutations in the *HBS1L* gene, who exhibited with growth restriction throughout intrauterine and postnatal life, developmental delay, and hypotonia (15, 16). The first ophthalmic examination at age 2 years disclosed no abnormalities. At 3 year of age the eyes were re-examined due to possible defective tracking, and abnormal retinal pigmentation was noted. At 4 years, her bilateral fundus photography revealed mottled pigmentation of the extramacular fundi at about the equator and 360 degrees (16). By 6 years, multiple, tiny half disc diameter hypopigmented spots with soft margins were noted in a broad equatorial band (*supplementary Figure S1*). There were rare pigment clumps among the hypopigmented lesions. Optical coherence tomography (OCT) images of each macula showed normal foveal architecture (*supplementary Figure S1*).

Electroretinography (ERG) was performed under anesthesia at age 7 years and showed signs of generalized retinal dysfunction and retinal dystrophy. Stimulus-response data were obtained in both the dark (scotopic) and light (photopic)-adapted conditions. Scotopic sensitivities (log sigma) were below the 99% prediction interval for normal (**Figure 1**) and saturated b-wave amplitudes (Vmax) were 92 μV (24%) and 80 μV (21%) of the normal mean, values that are below the 99% prediction interval for normal. The b-wave implicit times were prolonged. In light adapted photopic conditions, the amplitudes of the a- and b-wave responses were below the normal mean and implicit times were prolonged. (**Figure 2 and Figure 3**). Photopic function was also tested with 30Hz flickering white stimuli; response amplitudes were lower than normal (left eye: 46 μV, 37% of normal; right eye: 39 μV, 31% of normal).

**Figure 1:**
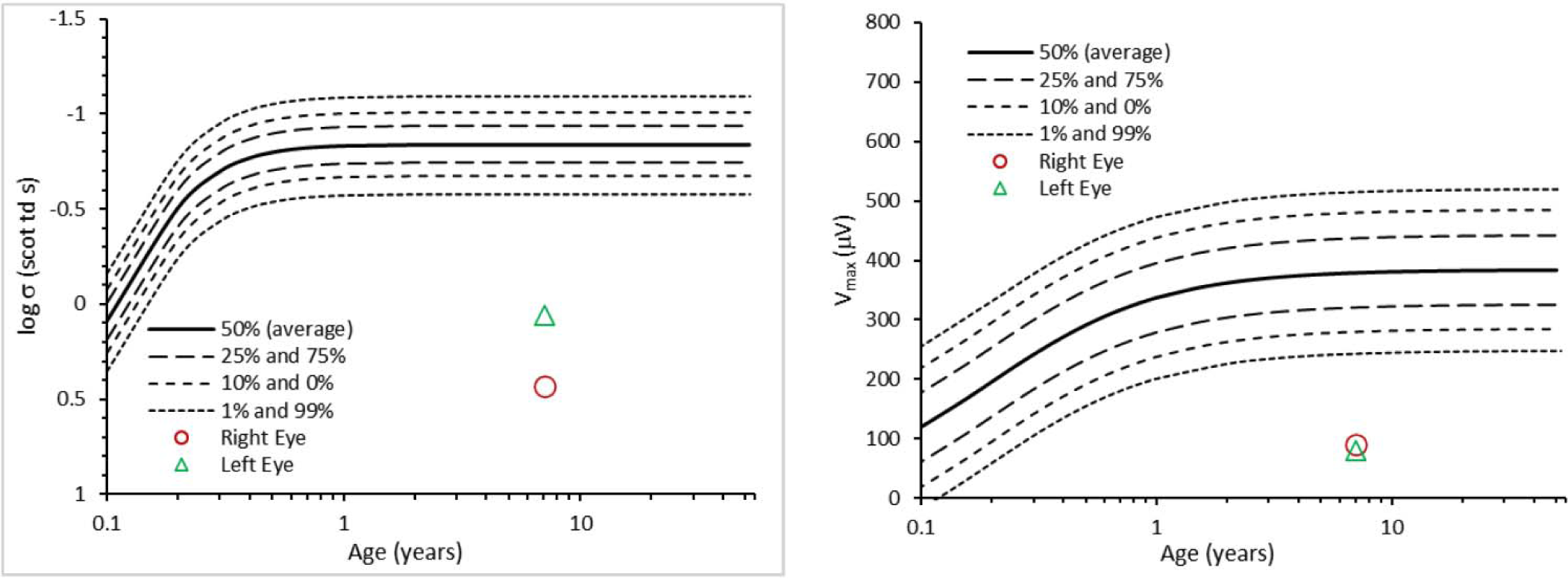
Electroretinography (ERG) shows remarkable affected scotopic function in the patient. Both log σ, an index of retina sensitivity, and the saturated b-wave amplitudes (Vmax) are below the 99% prediction interval for normal.

**Figure 2:**
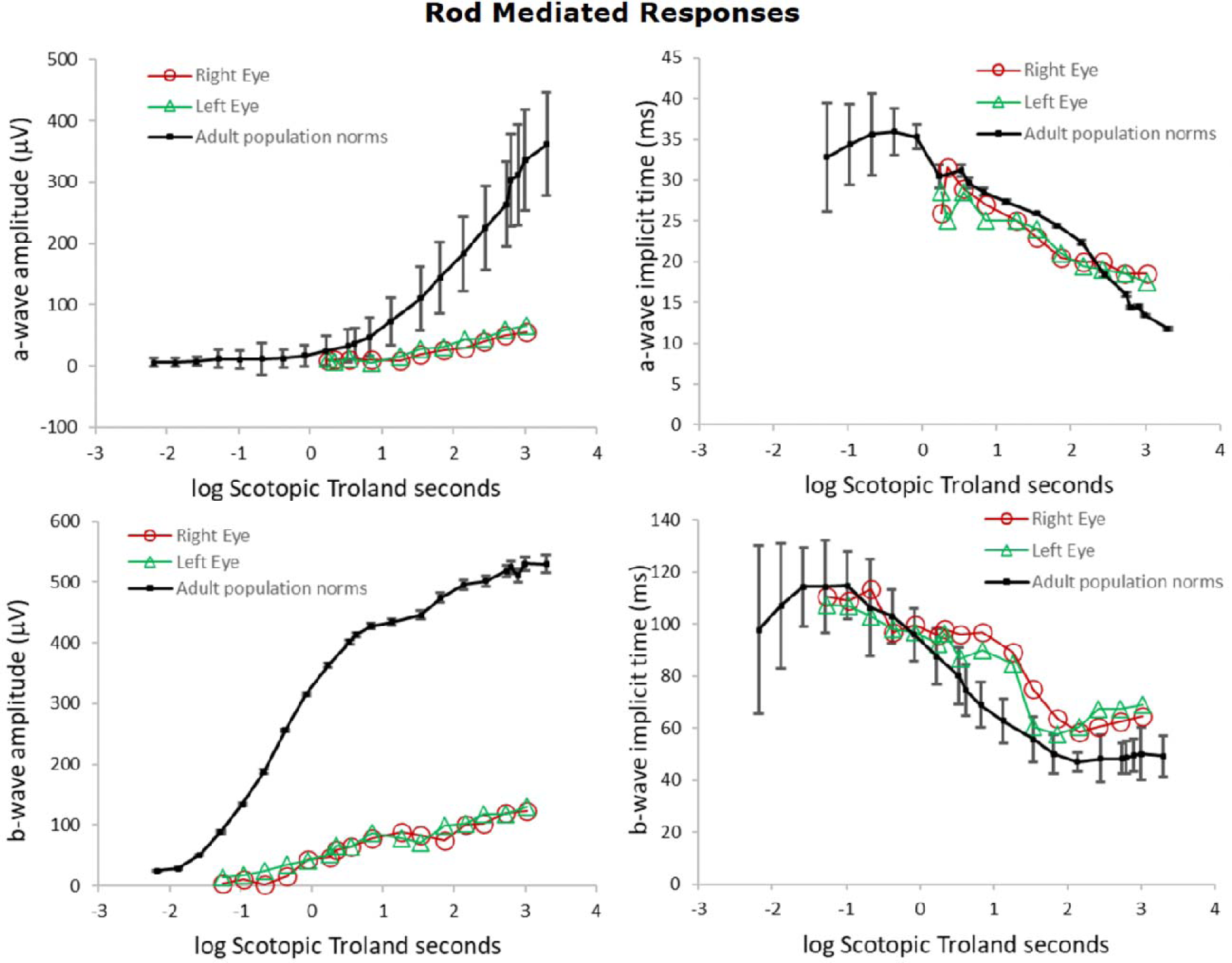
Electroretinography (ERG) shows severely damaged rod-mediated responses in the patient.

**Figure 3:**
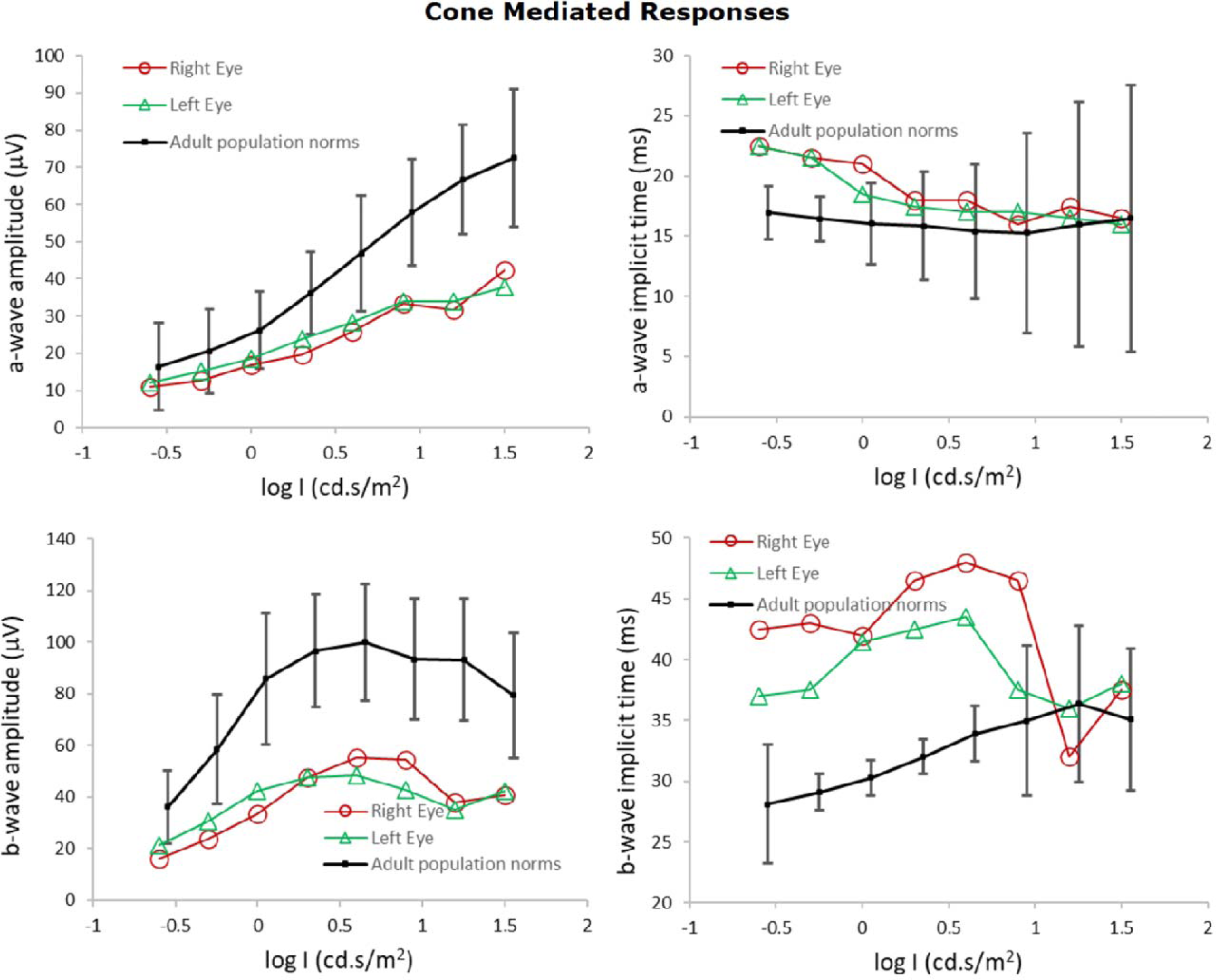
Electroretinography (ERG) shows impaired cone-mediated responses in the patient.

At age 7 years with correction of myopic astigmatism (−1.75 + 1.75 X 90 and −1.50 + 1.75 X 90) letter acuities were 20/50 for each eye. Dark adapted threshold (DAT) test was also performed using a 2 alternative forced choice psychophysical procedure which was within normal limits.

She has been followed until age 13 years. With slight up-dates in glasses prescriptions, acuities have been stable at 20/50, each eye. Fundus appearance has changed little over the last several years.

### Hbs1l deficiency causes retinal degeneration in *Hbs1l*^tm1a/tm1a^ mice

The RQC factors HBS1L and PELO are ubiquitously expressed in a variety of organs and tissues, such as brain, spinal cord, and skeletal muscles (16). To understand the transcript expression of *Hbs1l* in the retina, the web application eyeIntegration portal (https://eyeIntegration.nei.nih.gov) (25, 26) was used. The eyeIntegration incorporates single cell (sc) RNA-seq data from murine retina across two studies (27, 28), enabling visualization of single cell gene expression across multiple retinal cell types and different developmental time points, from embryonic day (E) 11 to postnatal day (P) 14. Our analysis of the data revealed that *Hbs1l* is broadly expressed in many types of cells during retinal development (*supplementary Figure S2*). At P8, its transcript level is relatively enriched in the photoreceptor and bipolar cells, and at P14, its expression appears to be decreased and restricted to some of the photoreceptor cells. This suggests that Hbs1l is broadly expressed during retinal development and enriched in photoreceptor cells.

A complete loss of Hbs1l leads to embryonic lethality (17), but marked reduction of *Hbs1l* (*Hbs1l*^tm1a/tm1a^) is associated with a viable mouse model with phenotypic overlap (16). The Hbs1l transcripts and proteins were markedly reduced in the retinas of these mice (*supplementary Figure S3*). The retinal structure and thickness was evaluated using optical coherence tomography (OCT) *in vivo* imaging in 4 week old mice. *Hbs1l*^tm1a/tm1a^ eyes showed decreased thickness of the whole retina compared with age-matched control mice (**Figure 4**, Mean ± S.D.: 134.0 ± 44.42 μm vs 173.1 ± 56.18 μm; *P* < 0.001), measured as the distance from the never fiber layer (NFL) to the retinal pigment epithelium (RPE), as well as decreased outer retina thickness (Mean ± S.D.: 55.56 ± 19.77 μm vs 93.92 ± 30.72 μm; *P* < 0.0001), measured from the outer plexiform layer (OPL) to the RPE. Histological analysis further confirmed that at 4 weeks of age, the average thickness of outer nuclear layer (ONL) was significantly reduced in the *Hbs1l*^tm1a/tm1a^ mice than the control group (**Figure 4**, Mean ± S.D.: 34.15 ± 9.28 μm vs 48.51 ± 13.56 μm; *P* < 0.0001). Consistently the outer segment (OS) and inner segment (IS) were also thinner in the *Hbs1l*^tm1a/tm1a^ mice compared to controls (**Figure 4**, Mean ± S.D.: 19.61 ± 5.43 μm vs 33.62 ± 9.22 μm; *P* < 0.0001). A similar structural pattern was observed in 2-week-old *Hbs1l*^tm1a/tm1a^ mice, with reduction in OS/IS thickness but less so of the ONL layer (*supplementary Figure S4*) compared to controls.

**Figure 4:**
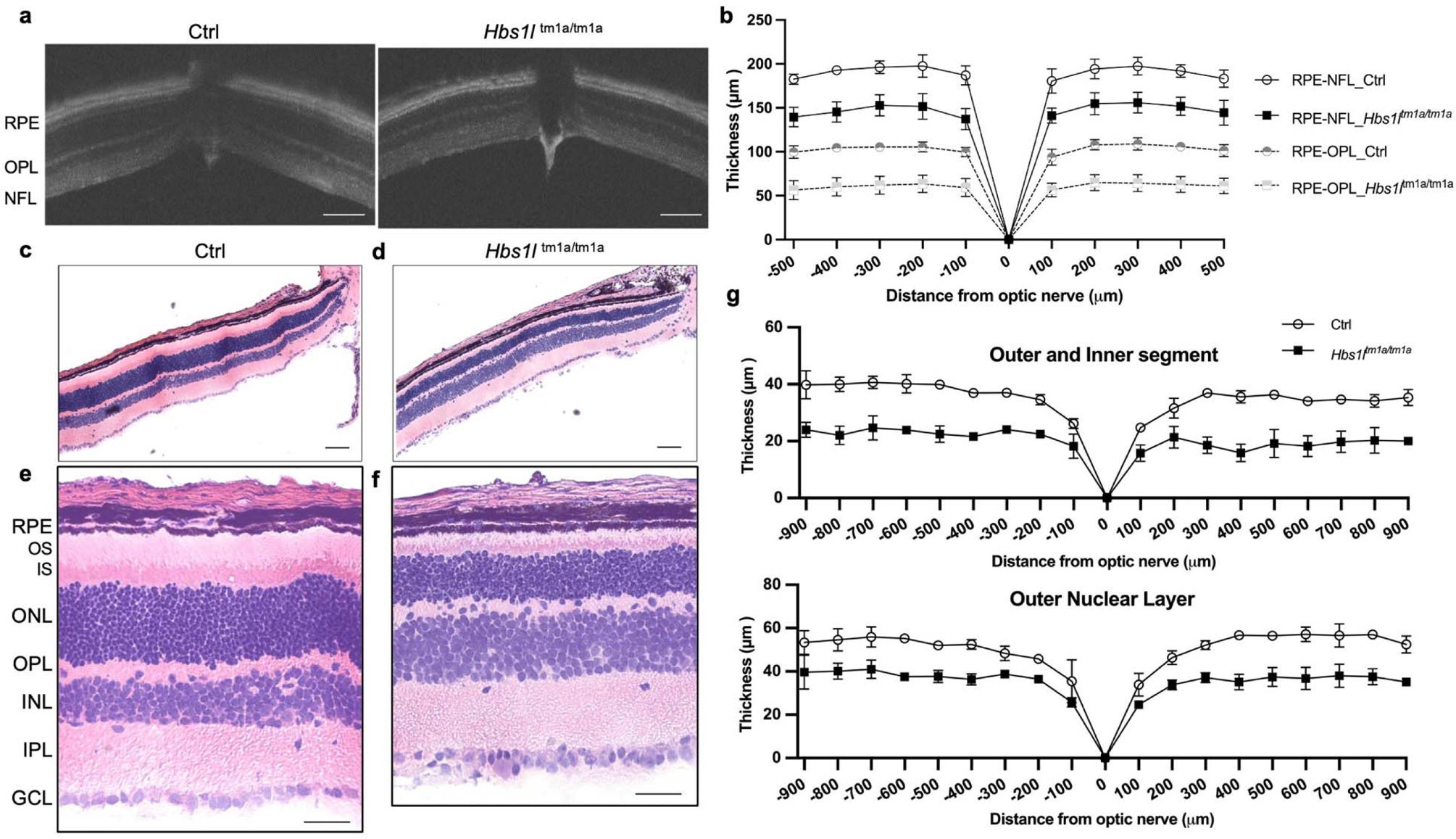
*Hbs1l*^tm1a/tm1a^ mice exhibited remarkable retinal dystrophy at 4 weeks of age. **(a)** Representative retinal images from optical coherence tomography (OCT) for thickness measurement and **(b)** its quantification. Thickness of both whole retina (RPE-NFL) and outer retina (RPE-OPL) were analyzed. Data are expressed as means ± S.D., n = 5 per group. \*\**P*<0.001 for all measurements (not shown in the figure). Scale bar = 100 μm. **(c-f)** Representative retinal images of H&E staining in the control (**c** and **e**) and *Hbs1l* ^tm1a/tm1a^ (**d** and **f**) mice. Panels **c** and **d** are retinal segments between optic nerve and approximately 1000 μm from the optic nerve with a scale bar of 100 μm. Panels **e** and **f** are retinal segments between 400 and 600 μm away from optic nerve with a scale bar of 100 μm. (**g**) Measurements of thickness of the outer segment (OS), inner segment (IS), and outer nuclear layer (ONL) in the control and *Hbs1l* ^tm1a/tm1a^ mice at 4 weeks of age. Data are shown as mean ± S.D., *n*=3 per group. Significant differences were calculated using nonparametric Mann-Whitney test. \*\*\**P*<0.0001. Ctrl, control; GCL, ganglion cell layer; INL, inner nuclear layer; IPL, inner plexiform layer; IS, inner segment; NFL, nerve fiber layer; ONL, outer nuclear layer; OPL, outer plexiform layer; OS, outer segment; RPE, retinal pigment epithelium; RPE-NFL, distance from RPE to NFL; RPE-OPL, distance from RPE to OPL.

To test if ONL thinning in the *Hbs1l*^tm1a/tm1a^ mice is associated with increased apoptosis of photoreceptor cells, we used terminal deoxynucleotidyl transferase (TdT)-mediated dUTP nick-end labeling (TUNEL) assay. At 2 weeks, the average number of apoptotic photoreceptor cells in the ONL of *Hbs1l*^tm1a/tm1a^ mice was significantly higher than that in the control group (Mean ± S.E.M.:105 ± 87 vs 17 ± 3, *P* = 0.0012), whereas the number of apoptotic cells in the INL did not change much in *Hbs1l*^tm1a/tm1a^ mice compared with control group (**Figure 5**). The ONL of *Hbs1l*^tm1a/tm1a^ mice at 4 weeks of age continue to show significant photoreceptor cell apoptosis but to a less extent (*supplementary Figure S5*). These results, along with the OCT and photoreceptor layer thickness analysis, suggested that loss of Hbs1l leads to premature death of photoreceptor cells and retinal thinning indicative of retinal degeneration.

**Figure 5:**
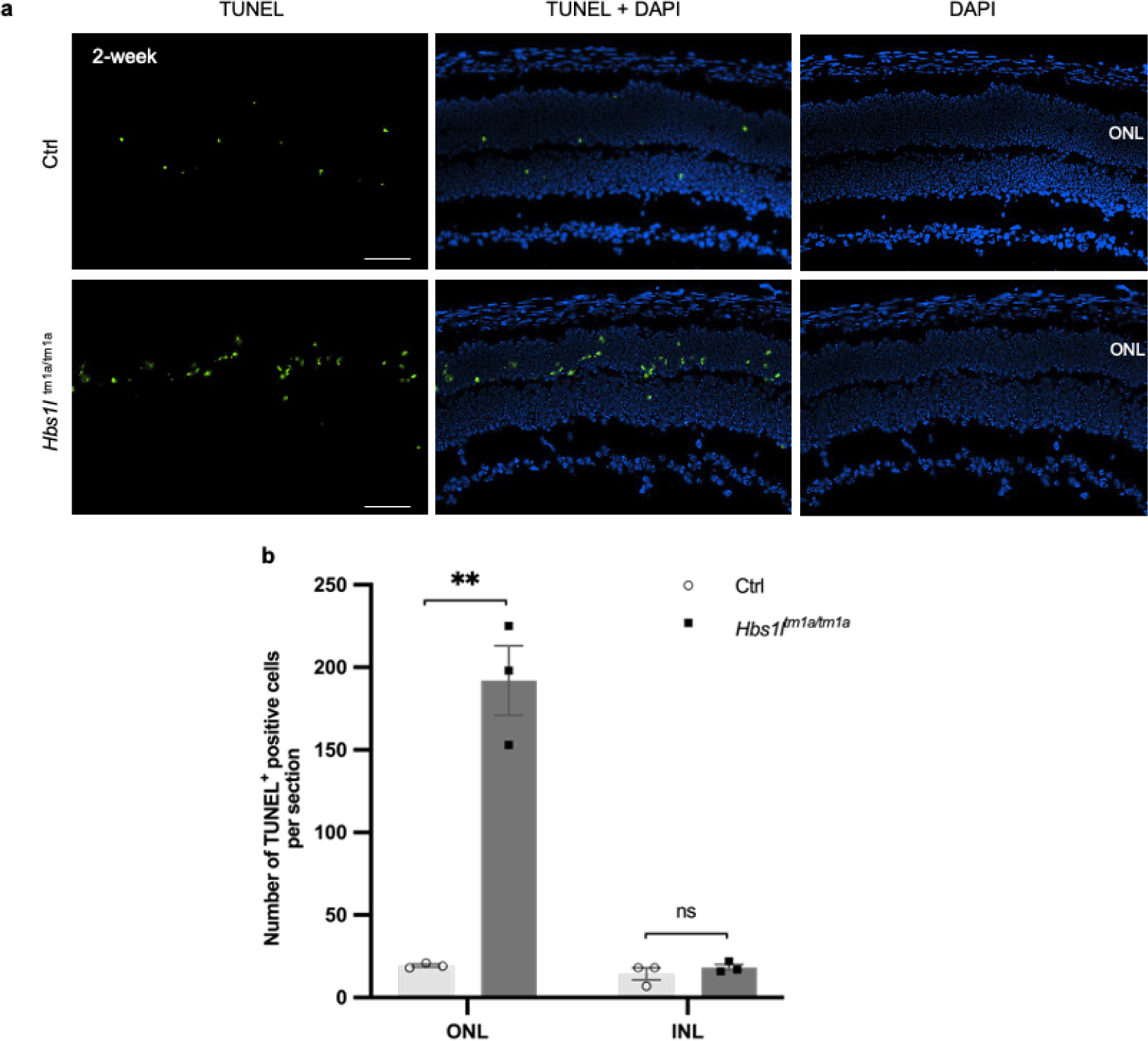
Loss of Hbs1l induces excessive apoptosis of photoreceptor cells. (**a**) Representative images of retinal sections from 2-week-old control (Hbs1l^+/+^) and *Hbs1l*^tm1a/tm1a^ mice. TUNEL (terminal deoxynucleotidyl transferase dUTP nick end labeling)-positive cells, green labelling; 4′,6-Diamidino-2-phenylindole (DAPI) counterstain for cell nuclei. Scale bar is 100 μm. (**b**) Histogram displaying the number of TUNEL-positive cells per section from 2-week-old control (Hbs1l^+/+^) and *Hbs1l*^tm1a/tm1a^ mice. Data represent mean ± S.E.M., *n*=3 per group. Two-tailed Student’s t-test was used for statistical analysis. \*\**P* < 0.01. Ctrl, control; ONL, outer nuclear layer; INL, inner nuclear layer.

### Quantitative profile analysis of retinal proteome in the *Hbs1l*^tm1a/tm1a^ mice using multiplexed TMT

Loss of Hbs1l is known to alter the transcription regulation and induce defects in translation elongation (17), although its resultant proteomic changes haven’t been deciphered in the retinas. In this study, quantitative mass spectrometry (MS) analysis of retinal proteome in *Hbs1l*^tm1a/tm1a^ mice was performed and compared to littermate controls. A total of 8114 proteins from retinal tissue were quantified by multiple peptides at an initial protein false discovery rate (FDR) of less than 1% (**Figure 6a**, see *supplementary Table S1*). The quality of the data and reproducibility of the biological replicates across groups were assessed using statistical metrics, hierarchical clustering and PCA analysis (**Figure 6b** and *supplementary Figure S6*).

**Figure 6:**
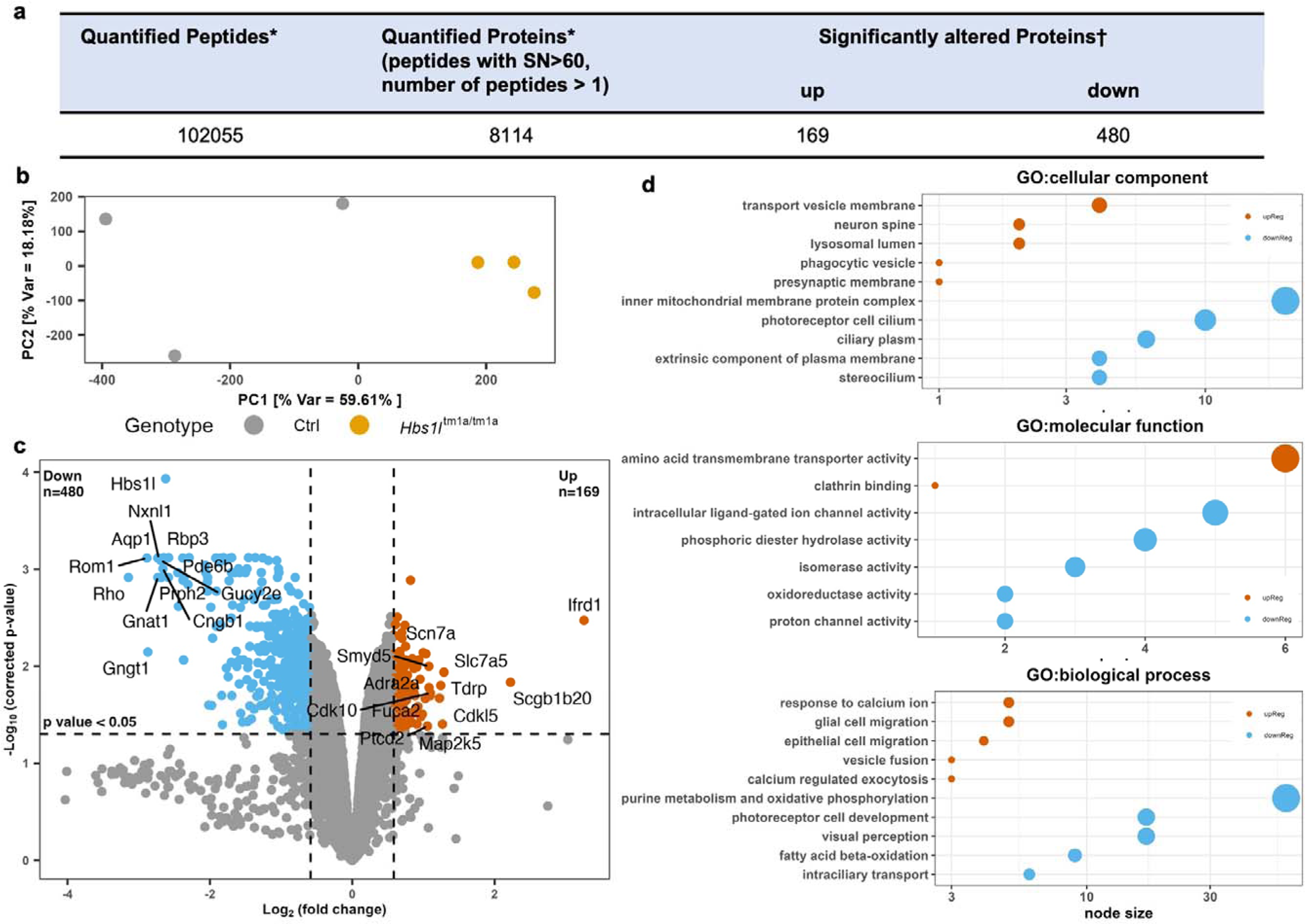
Summary of retinal proteomics data set and Gene Ontology (GO) analysis of significantly enriched proteins in *Hbs1l*^tm1a/tm1a^ and control mice. **(a)** Table of quantified peptides and proteins in this experimental data set. Asterisks indicate quantified across all 6 channels with SN>60 and number of peptides>1. Daggers indicate a Benjamini−Hochberg-corrected pvalue of <0.05 and a fold change beyond ±1.5. **(b)** Principal component analysis of the biological replicates (*n*=3 per group). **(c)** Volcano plot displaying the −log_10_ (corrected p value) vs log_2_ (fold change) for all quantified proteins. **(d).** A total of 649 DMPs (169 up regulated and 480 down regulated) were used for GO analysis using g:Profiler (https://biit.cs.ut.ee/gprofiler). An enrichment map was created with parameters FDR Q value < 0.01, and combined coefficient >0.375 with combined constant = 0.5. Clusters of nodes were labeled using the AutoAnnotate Cytoscape application. The top 5 GO clusters were shown for illustration; only 2 clusters were formed for GO: molecular function from upregulated proteins. The size of filled circles indicates the size of each cluster of nodes.

Student t test was used to determine proteins that are significantly altered between the *Hbs1l*^tm1a/tm1a^ and control mice. Proteins that met the p-value cut-off (≤0.05) and at least 50% difference in levels (*supplementary Table S2*) were considered as differentially modulated proteins (DMPs). This two-step differential analysis of *Hbs1l*^tm1a/tm1a^ versus control for retina tissue yielded 169 proteins with increased level (p-value_≤_0.05 and_≥_1.5-fold change) and 480 decreased proteins (p-value_≤_0.05 and ≤ 0.67-fold change) (**Figure 6c**). Hierarchical clustering analysis of proteins with differential abundance illustrated the overall consistency of the up and down regulation within respective *Hbs1l*^tm1a/tm1a^ and control group (*supplementary Figure S6*).

Following this, to examine the biological functions of DMPs, we performed enrichment analysis of GO and reactome pathway (*supplementary Table S3)* using g:Profiler (https://biit.cs.ut.ee/gprofiler). The DMPs were significantly (p_<_0.01) enriched in the inner mitochondrial membrane protein complex, photoreceptor cell cilium, and transport vesicle membrane, which are involved in the molecular functions related to oxidoreductase and proton channel activity, intracellular ligand-gated ion channel activity, and amino acid transmembrane transporter activity (**Figure 6d** and *supplementary Figure S7*). Furthermore, GO biological process analysis revealed that the DEPs in *Hbs1l*^tm1a/tm1a^ mice were mainly involved in processes related to purine metabolism and oxidative phosphorylation, photoreceptor cell development and visual perception, which were also identified by Reactome pathway analysis (*supplementary Figure S8*).

To explore the comprehensive effects of Hbs1l deficiency in the retina, we performed unbiased gene set enrichment analysis (GSEA) using the retinal proteomics data. Among 7775 gene sets categorized to gene ontology biological process, 2715 gene sets were used in the analysis after gene set size filtering (min=15, max=250), of which 383 gene sets were altered with statistical significance (FDR q value less than 0.05) and visualized with EnrichmentMap from Cytoscape (*supplementary Figure S9*). The size of a node indicates the size of each gene set, and the edge means two connected nodes share some genes. In accordance with GO and reactome pathway analysis using g:Profiler, gene sets related to aerobic respiration, photoreceptor differentiation, phototransduction, and cilium assembly were reduced in the *Hbs1l*^tm1a/tm1a^ mice while gene sets related to neurotransmission, synapse assembly, axonogenesis, and mononuclear cell migration were enriched in Hbs1l-deficient retina.

### Validation of differentially modulated proteins (DMPs) using Western Blot

Several proteins that participate in the phototransduction cascade of rod photoreceptors, such as rhodopsin (Rho), transducing subunits (Gngt1, Gnat1 and Gnb3), cGMP phosphodiesterase subunits (Pde6a and Pde6b), and cyclic nucleotide gated (CNG) channel structure (Cngb1, Cnga1), were significantly reduced (**Figure 6c** and *supplementary Table S2*). Other decreased proteins included structural proteins essential for photoreceptor disc morphogenesis such as Rom1. Interestingly, mutations in Rho, Pde6a/Pde6b and Rom1 are associated with inherited retinal degeneration in both human and mice, and their downregulation may underlie the retinal degenerative phenotype as observed in *Hbs1l*^tm1a/tm1a^ mice. On the other hand, the expression of apoptosis associated proteins including Ddit3, Dapl1 and Dap were increased in the *Hbs1l*^tm1a/tm1a^ mouse retina.

Western blot (**Figure 7**) confirmed the remarkable decrease of Rho protein expression but not Opn1sw, a cone opsin for color vision, indicating that the observed retinal degeneration affects mostly rods but not cones at this time point. Consistent with our previous findings (16), loss of Hbs1l is accompanied by decreased levels of its partner protein Pelo while the protein level of Abce1 remain unchanged. Interestingly, after searching for any deregulated ribosome-related proteins besides Pelo, we found decreased levels of translation machinery associated 7 homolog (Tma7) and Endothelial differentiation-related factor 1(Edf1) in Hbs1l-deficient retina. Tma7 is a ubiquitously expressed protein that may inhibit autophagy by activating the PI3K/mTOR pathway (22), and Edf1 is reported to coordinate cellular responses to ribosome collisions (23, 24). In addition to rescuing the ribosomes stalled at the mRNA 3’ end (29), it remains to be elucidated if HBS1L/PELO may play a role in resolving collided ribosomes.

**Figure 7.**
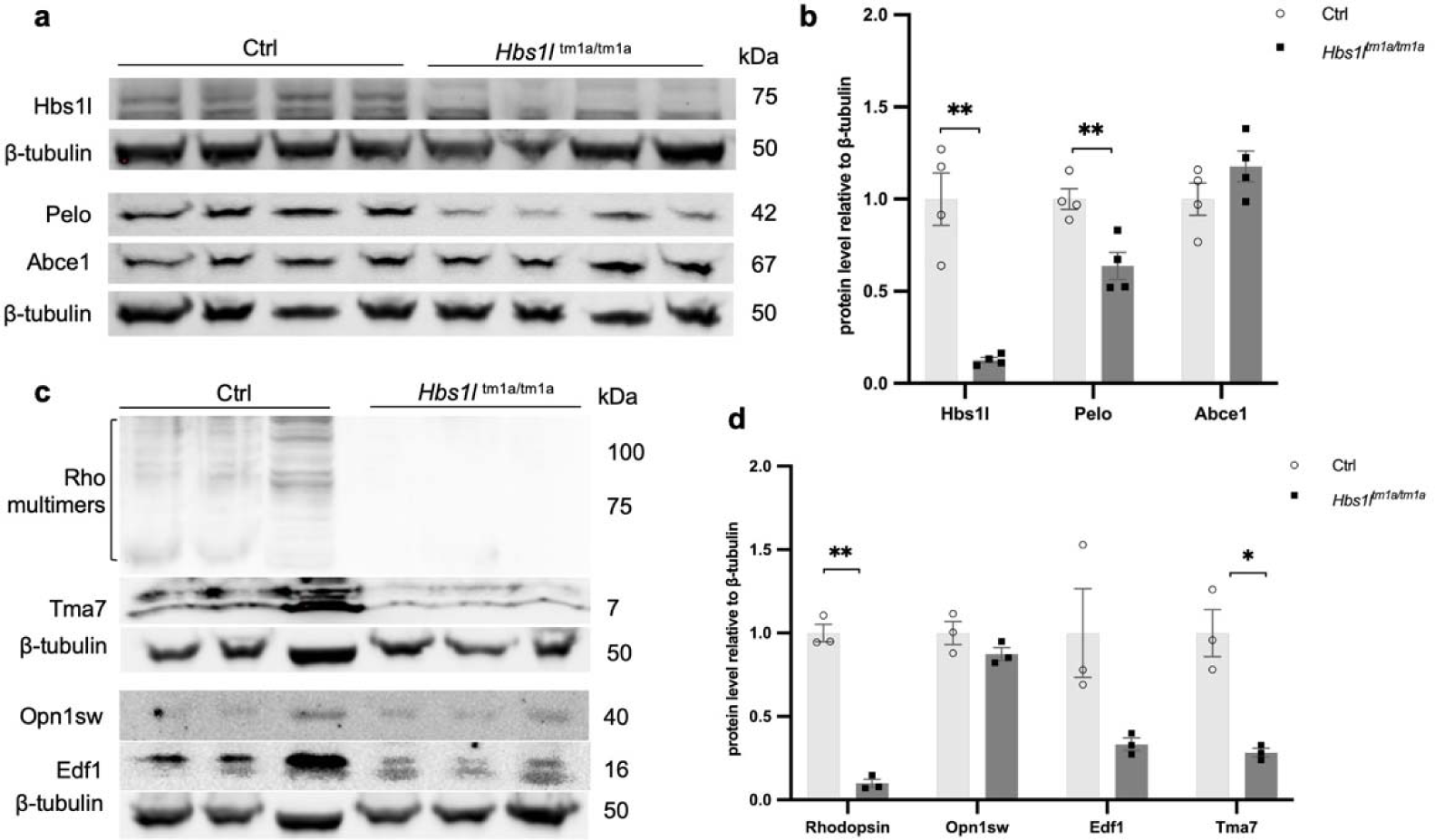
Hbs1l deficiency leads to a reduction in the protein levels of Pelo, Tma7, and Edf1. along with many retinal proteins critical for visual cascade (Rho as an example). Representative western blot images **(a** and **c)** and quantification (**b** and **d**) by ImageJ analyses. Data represent mean ± S.E.M., *n* = 3-4 per group, \**P* < 0.05, \*\**P* < 0.01.

## Discussion

In this study, we report that loss of HBS1L results in retinal degeneration in both human patient and the *Hbs1l*^tm1a/tm1a^ hypomorph mouse model. OCT and H&E staining analyses distinctly indicate that the thickness of total retina, outer nuclear layer (ONL), outer segment (OS) and inner segment (IS) were decreased in *Hbs1l*^tm1a/tm1a^ mice as early as 2 weeks of age. At this time, the thickness of OS/IS layer was dramatically decreased even when normalized by the total retinal thickness. This reduction correlates with an increased apoptosis in the ONL as evidence by the TUNEL assay. Consistent with the pathological phenotype observed in *Hbs1l*^tm1a/tm1a^ mice, proteomic analysis revealed a significant decrease in the levels of many critical retinal proteins, including photoreceptor visual signaling protein (such as Rho) and structural proteins (Rom1), along with elevated levels of apoptosis-related proteins.

The findings of OS/IS thinning at 2 weeks of age suggests that OS was probably not well formed although increased apoptosis suggests degenerative process playing a role. We hypothesize that Hbs1l deficiency leads to impaired production of pan retinal proteins, especially those required for proper disc morphogenesis. Due to this blunted rod OS disc morphogenesis as well as an imbalance between protein synthesis and degradation, it sets the stage for the photoreceptor cell apoptosis and retinal degeneration. In the meantime, without Hbs1l, the accumulation of dysfunctional ribosomes/proteins may overwhelm the lysosomes and autophagy pathways for protein degradation, thereby exaggerating the cell apoptosis.

Historically yeast Hbs1 and its mammalian counterpart HBS1L have been recognized for their role as a ribosomal rescue factor and play critical roles in processes of no-go decay (13, 30), non-stop decay (31, 32), dissociation of vacant or stress-related non-translating inactive 80S ribosomes (12, 33), as well as in maintaining oxidative stress resilience (34) and endoplasmic reticulum homeostasis (35). The absence of HBS1L is accompanied by a reduction in the protein level of its interacting partner PELO (16), which contributes to an elevated baseline of 80S monosomes (16) and ribosomal pausing during transcription elongation (17). The intriguing aspect is why decreased protein expression of ribosomal rescue proteins, like HBS1L, elicits tissue-specific phenotypes. While *Hbs1l*^tm1a/tm1a^ hypomorph mice do not display overt neurodegenerative phenotype (17), likely due to residual protein expression, an equivalent reduction of Hbs1l leads to retinal dystrophy in these mice. This difference may be partially explained by the extraordinary demand of protein synthesis for disc membrane and outer segment renewal in the retina (36–38) along with limited actively translating ribosomes due to Hbs1l depletion (16).

Cryo-electron microscopic (EM) analysis of Dom34 (paralog of Pelo in yeast) bound to a ribosome stalled at the mRNA 3’ end has yielded novel insights into how the sensing of such stalled ribosomes occurs (29). The N-terminal domain of Hbs1 binds to the ribosome near the entry to the mRNA channel, whereas Dom34 binds to an empty ribosomal A site, inserting a β loop into the channel. Once Dom34/PELO is bound, Hbs1/HBS1L dissociates, and Dom34/PELO recruits the ATPase Rli1/ABCE1 which prompts the splitting of the two ribosomal subunits. Despite their efficient recycling of stalled ribosome at the mRNA 3’ end, the requirement for PELO to protrude into the decoding center renders them incapable of splitting internally stalled ribosomes during translation. Under such circumstances, a distinct recycling pathway (ZNF598 and ASC-1) is necessary for disassemble collided ribosomes (39–41).

In this study, we discovered that the protein level of Edf1 and Tma7 were markedly diminished following HBS1L deletion. EDF1, initially recognized for its role in endothelial function (42), has recently been identified as a novel protein recruited to collided ribosomes during translational distress (23, 24). While the loss of EDF1 leads to a reduction in ZNF598-mediated ubiquitylation of ribosomal proteins, the recruitment of EDF1 to collided ribosomes occurs independently of ZNF598. Cryo-EM analyses of EDF1 and its yeast counterpart Mbf1 have revealed a conserved binding site on the 40S ribosomal subunit at the mRNA entry channel near the collision interface. Furthermore, mapping the EDF1 interactome revealed translational repressors GIGYF2•EIF4E2, along with known components of the RQC machinery including HBS1L. It remains to be elucidated whether HBS1L may interact with EDF1 and participate in the rescue of internal stalls, especially ASCC-resistant stalls (43), as well as how the loss of HBS1L/PELO may affect the level of these proteins.

In summary, we reported that genetic deficiency of HBS1L causes retinal degeneration in both the human patient and *Hbs1l*^tm1a/tm1a^ mice. We also present a comprehensive profile of the proteomic changes in the *Hbs1l*^tm1a/tm1a^ retina, revealing the underlying disruption of disc morphogenesis and photoreceptor cell development. Moreover, the loss of Hbs1l results in reduced levels of Pelo, Edf1, and Tma7 proteins, although their intricate ribosomal regulatory mechanisms warrant further exploration.

## Materials and methods

### Ethics statement

The IRB at Boston Children’s Hospital approved the human subject study under the protocol 10-02-0253. The Institutional Animal Care and Use Committee (IACUC) at Boston Children’s Hospital approved our mouse work (approval number 16-06-3182R). The work followed the Guide for the Care and Use of Laboratory Animals and all of the regulatory protocols set forth by the Boston Children’s Hospital Animal Resources at Children’s Hospital (ARCH) facility.

### Mouse strain and genotyping

The “Knockout-first” *Hbs1l*^tm1a^ (C57BL/6N-*A^tm1Brd^Hbs1l^tm1a(KOMP)Wtsi^*, MMRRC:048037) mice were obtained from the International Knockout Mouse Consortium (Knockout Mouse Project (KOMP) Repository, IKMC project 79564; http://www.mousephenotype.org/data/alleles/MGI:1891704/tm1a(KOMP)Wtsi), which is developed by the Wellcome Trust Sanger Institute (Hinxton CB10 1SA, UK) (44). Mice were quarantined at Charles River Laboratories (Worcester, MA, USA) before import into our animal facility at Boston Children’s Hospital (Boston, MA, USA). All experiments and quantifications were performed with at least three mice of each genotype and time point using mice of either sex. Absence of Crb1 (rd8) mutation was confirmed by PCR to rule out potential confounding effects by rd8 (45).

Genotypes were confirmed by PCR using the *Hbs1l* forward primer (Hbs1l_F: 5’- TCTAATTCATGTGTGCCGCC-3’) and reverse primer (Hbs1l_R: 5’- TCCTGTGTTTTACCTGCATAGAGC’3’) which flanked the targeted exon 5, producing the WT PCR product of 483 bp. Addition of the targeting cassette specific primer (Hbs1l_Cas: 5’- TCGTGGTATCGTTATGCGCC-3’) to the PCR master mixes to induce multiplex reactions generated a mutant-specific allele of 338 bp.

### Reverse transcription and quantitative PCR (qPCR) analysis

Adult mouse eye tissues were isolated and immediately frozen in liquid nitrogen. Total RNA was extracted with Trizol reagent (Life Technologies). cDNA synthesis was performed on DNase-treated (DNA-free DNA Removal Kit, Life Technologies AM1906) total RNA using oligo(dT) primers and SuperScript III First-Strand Synthesis System (Life Technologies). Quantitative RT-PCR reactions were performed using iQ SYBR Green Supermix (Bio-Rad) and an CFX96 Real-Time PCR Detection System (Bio-Rad). Expression levels of β*-actin* were used as input control for semi-quantitative RT-PCR. For quantitative RT-PCR (qPCR) analysis, expression levels of the genes of interest were normalized to β*-actin* using the 2^-ΔΔCT^ method (46) and expressed as the fold change + standard error of the mean (SEM) relative to control.

Semi-quantitative RT-PCR Primers (F, Forward; R, Reverse):

- *Hbs1l Exon 3* F: 5’GAAATTGACCAAGCTCGCCTGTA3’
- *Hbs1l Exon 6* R: 5’CTCAGAAGTTAAGCCAGGCACT3’
- β*-actin F:* 5’AGGCCAACCGTGAAAAGATG3’
- β*-actin* R: 5’AGAGCATAGCCCTCGTAGATGG3’

Quantitative RT-PCR Primers (F, Forward; R, Reverse):

- *Hbs1l* F: 5’AGACCATGGGATTTGAAGTGC3’
- *Hbs1l* R: 5’CCGGTCTCAGGAATGTTAGGA3’
- *Hbs1l II* F: 5’TGAAGTTGAACAAAGTGCCAAG3’
- *Hbs1l II* R: 5’CTGCTTCCTCTGTGTTCCTC3’
- β*-actin F:* 5’AGGCCAACCGTGAAAAGATG3’
- β*-actin* R: 5’AGAGCATAGCCCTCGTAGATGG3’

### Optical coherence tomography (OCT) imaging

OCT imaging was performed as previously described (47, 48). In brief, mice were anesthetized with a mixture of xylazine (6 mg/kg) and ketamine (100 mg/kg), and pupils were dilated with topical drops of Cyclomydril (Alcon Laboratories, Fort Worth, TX). Two minutes after pupil dilation, lubricating eye drops (Alcon Laboratories) were applied to the cornea. Spectral domain OCT with guidance of bright-field live fundus image was performed using the image-guided OCT system (Micron IV, Phoenix Research Laboratories) according to the manufacturer’s instruction and using the vendor’s image acquisition software to generate bright field images, and OCT scans.

### Histology, immunofluorescence, and TUNEL assay

The mouse eyeballs were enucleated, embedded in OCT Tissue Tek Compound (Sakura Finetek, USA), and placed in isopentane cooled by liquid nitrogen. Hematoxylin and eosin staining was carried out on 5_μm thickness cryosections. Immunofluorescence was performed by standard protocol using rabbit anti-HBS1L polyclonal antibody (10359-1-AP, 1:50 dilution, Proteintech Group Inc., Chicago, IL, USA) and rabbit anti-PELO polyclonal antibody (10582-1-AP, 1:100 dilution, Proteintech Group Inc.). TUNEL staining was performed using the TACS TdT In Situ Apoptosis Detection Kit (Trevigen, Gaithersburg, MD, USA) on retinal cryosections. Staining was carried out according to the manufacturer’s instructions. The nuclei were counterstained with 4′,6-Diamidino-2-phenylindole (DAPI). Images were captured using a Nikon Eclipse 90i microscope in conjunction with NIS-Elements AR software (Nikon Instruments Inc., NY, USA). Six fields of view per section at 20× magnification was evaluated to determine the average number of apoptotic cells.

### Mass spectrometry using Tandem Mass Tag (TMT)

The retinal tissues from three pairs of *Hbs1l*^tm1a/tm1a^ mice and littermate controls at 4 weeks of age were isolated and mass spectrometry was performed by the Thermo Fisher Scientific Center for Multiplexed Proteomics (TCMP) at Harvard Medical School, as described previously (49). Briefly, tissues are lysed by bead-beating and protein are quantified using the Pierce BCA Protein Assay Kit (Thermo-Scientific). Sample were reduced with tris(2-carboxyethyl)phosphine (TCEP), alkylated with Iodoacetimide, quenched with DTT. Protein was chloroform - methanol precipitated, reconstituted, and digestion was performed sequentially using LysC (1:50) and trypsin (1:100) based on protease to protein ratio. 6-Plex TMT labeled peptides from 6 samples were mixed into one sample, desalted and fractionated with basic-pH reverse-phase (BPRP) high-performance liquid chromatography (HPLC), collected in a 96-well plate and combined to make a set of 12 fractions for liquid chromatography−tandem mass spectrometry (LC−MS/MS) processing.

Mass spectrometric data were collected on an Orbitrap Eclipse™ Tribrid™ Mass Spectrometer (ThermoFisher Scientific). MS/MS spectra are matched to a protein database using a SEQUEST-based in-house built software platform (49). MS2 spectra were searched using the COMET algorithm (using a peptide mass tolerance of 50 ppm and fragment ion tolerance of 1.005 Da) against a Mouse Uniprot composite database containing its reversed complement and known contaminants. For whole proteome analysis, only Methionine oxidation is used as differential modification allowing up to two internal cleavage sites and up to three differential sites. Peptide spectral matches were filtered to a 1% false discovery rate (FDR) using the target-decoy strategy combined with linear discriminant analysis. The proteins were filtered to a <1% FDR and quantified only from peptides with a summed SN threshold of >60.

To control for differential protein loading within a six-plex, the summed protein quantities were adjusted to be equal within each channel. Following this, relative protein abundance was calculated as the ratio of sample abundance to total abundance using the summed reporter ion intensities from peptides that could be uniquely mapped to a gene. Small differences in laboratory conditions and sample handling can result in systematic, sample-specific bias in the quantification of protein levels. To mitigate these effects, the relative abundances were log2 transformed and zero-centered at the protein level to obtain final relative abundance values, and then we computed the median, log2 transformed relative protein abundance for each sample and re-centered to achieve a common median.

### Bioinformatics and functional pathway analysis

Differentially expressed proteins were assessed using a two-sample t test (P_≤_0.05) and a fold change threshold (≥1.5 for upregulation or ≤0.67 for downregulation). The reproducibility of all experiment output was evaluated by further statistical analysis such as overall data quality, un-supervised analyses such as clustering and principal component analysis (PCA). All data analyses were performed using R (v4.2.1; R Core Team 2022) (50) and RStudio (v2022.7.1.554; RStudio Team 2022) (51) statistical software unless otherwise stated. Some specific libraries used for plotting include cowplot (v1.1.1; Wilke C 2020) (52), ggplot2 (v3.4.0; H. Wickham 2016) (53), ggrepel (v0.9.2; Slowikowski K 2022) (54), gplots (v3.1.3; Warnes G et al. 2022) (55), and dendextend (v1.16.0; Tal Galili 2015) (56). Network graphics were generated in Cytoscape (v3.9.1, National Institute of General Medical Sciences, Bethesda, USA) (57).

The GO of proteins was classified using g:Profiler (https://biit.cs.ut.ee/gprofiler)(58) to explore the functionality of altered GO biological process and reactome pathways in Hbs1l-deficient retina. A cutoff of p value < 0.05 was adjusted for all GO categories. Gene Set Enrichment Analysis (GSEA) was performed using GSEA software (59, 60) and Molecular signatures database (MSigDB) (61, 62). GSEA enrichment analysis results were reduced and summarized by using the EnrichmentMap Pipeline Collection (v1.1.0) from Cytoscape (63–66).

### Western blot

The retinal tissues from *Hbs1l*^tm1a/tm1a^ mice and littermate controls were dissected, snap frozen in isopentane, and stored at −80°C until analysis. Protein isolation and western blot procedures were performed as described previously (67). Proteins were probed with antibody against HBS1L (10359-1-AP, Proteintech, 1:500 dilution), PELO (10582-1-AP, Proteintech, 1:500 dilution), ABCE1 (ab32270, Abcam, 1:1,000 dilution), Rhodopsin (MABN15, Millipore, 1:500 dilution), Opn1sw (sc-14365, Santa Cruz, 1:500 dilution), TMA7 (20393-1-AP, Proteintech, 1:500 dilution), and EDF1 (12419-1-AP, Proteintech, 1:500 dilution). Quantification of protein levels normalized to β-tubulin (12004165, Bio-Rad, 1:4000 dilution) was performed using Image J software.

### Statistical analysis

Data were analyzed with GraphPad Prism (v.9.0; GraphPad Software) and reported as mean ± standard deviation (S.D.) or mean ± standard error of the mean (S.E.M.). Unpaired two-tailed Student’s t-test was used for statistical analysis unless indicated otherwise. Figure 4 used nonparametric Mann-Whitney test for group comparison of retinal layer thickness. *P* value of less than 0.05 were considered significant. The numbers of samples per group (n) and statistical significance for all comparisons are specified in the figure legends.

## Data availability

The mass spectrometry proteomics data have been deposited to the ProteomeXchange Consortium via the PRIDE partner repository (68–70) with the dataset identifier PXD045660. The complete dataset including the complete list of proteins identified can be found in Supplementary Table S1.

## Acknowledgement

The authors would like to thank the patient and her parents for their commitment to this research and to finding a cause and a cure for this disorder. In addition, we thank the Wellcome Sanger Institute Mouse Genetics Project (Sanger MGP) and its funders for providing the mutant mouse line (*Hbs1l*^tm1a/tm1a^). We also thank the Rodent Histopathology Core and the Thermo Fisher Scientific Center for Multiplexed Proteomics at Harvard Medical School (http://tcmp.hms.edu) for their technical support. KB, AKB and JC were supported by NIH R01 grants (EY024963, EY028100, EY 031765, to JC).

## Author contributions

SL and PBA designed the experiments and project administration. All authors participated in performing the experiments, data analyses, manuscript drafting and/or figure generation. All the authors read and approved the final manuscript.

## Declaration of interests

The authors declare no financial or competing interests.

## Notes

### Competing Interest Statement

The authors have declared no competing interest.

## Reference

1. Jamar NH, Kritsiligkou P, Grant CM. Loss of mRNA surveillance pathways results in widespread protein aggregation. Scientific Reports. 2018;8(1).

2. Brandman O, Hegde RS. Ribosome-associated protein quality control. Nature Structural &amp; Molecular Biology. 2016;23(1):7–15.

3. Graille M, Séraphin B. Surveillance pathways rescuing eukaryotic ribosomes lost in translation. Nat Rev Mol Cell Biol. 2012;13(11):727–35.

4. Shoemaker CJ, Green R. Translation drives mRNA quality control. Nature Structural &amp; Molecular Biology. 2012;19(6):594–601.

5. Joazeiro CAP. Mechanisms and functions of ribosome-associated protein quality control. Nature Reviews Molecular Cell Biology. 2019;20(6):368–83.

6. Shao S, von der Malsburg K, Hegde RS. Listerin-dependent nascent protein ubiquitination relies on ribosome subunit dissociation. Mol Cell. 2013;50(5):637–48.

7. Choe Y-J, Park S-H, Hassemer T, Körner R, Vincenz-Donnelly L, Hayer-Hartl M, et al. Failure of RQC machinery causes protein aggregation and proteotoxic stress. Nature. 2016;531(7593):191–5.

8. Ishimura R, Nagy G, Dotu I, Zhou H, Yang X-L, Schimmel P, et al. Ribosome stalling induced by mutation of a CNS-specific tRNA causes neurodegeneration. Science. 2014;345(6195):455–9.

9. Yang J, Hao X, Cao X, Liu B, Nystrom T. Spatial sequestration and detoxification of Huntingtin by the ribosome quality control complex. Elife. 2016;5.

10. Rimal S, Li Y, Vartak R, Geng J, Tantray I, Li S, et al. Inefficient quality control of ribosome stalling during APP synthesis generates CAT-tailed species that precipitate hallmarks of Alzheimer’s disease. Acta Neuropathol Commun. 2021;9(1):169.

11. Nelson RJ, Ziegelhoffer T, Nicolet C, Werner-Washburne M, Craig EA. The translation machinery and 70 kd heat shock protein cooperate in protein synthesis. Cell. 1992;71(1):97–105.

12. van den Elzen AM, Schuller A, Green R, Séraphin B. Dom34-Hbs1 mediated dissociation of inactive 80S ribosomes promotes restart of translation after stress. Embo j. 2014;33(3):265–76.

13. Shoemaker CJ, Eyler DE, Green R. Dom34:Hbs1 promotes subunit dissociation and peptidyl-tRNA drop-off to initiate no-go decay. Science. 2010;330(6002):369–72.

14. Kalisiak K, Kuliński TM, Tomecki R, Cysewski D, Pietras Z, Chlebowski A, et al. A short splicing isoform of HBS1L links the cytoplasmic exosome and SKI complexes in humans. Nucleic Acids Res. 2017;45(4):2068–80.

15. Sankaran VG, Joshi M, Agrawal A, Schmitz-Abe K, Towne MC, Marinakis N, et al. Rare complete loss of function provides insight into a pleiotropic genome-wide association study locus. Blood. 2013;122(23):3845–7.

16. O’Connell AE, Gerashchenko MV, O’Donohue MF, Rosen SM, Huntzinger E, Gleeson D, et al. Mammalian Hbs1L deficiency causes congenital anomalies and developmental delay associated with Pelota depletion and 80S monosome accumulation. PLoS Genet. 2019;15(2):e1007917.

17. Terrey M, Adamson SI, Chuang JH, Ackerman SL. Defects in translation-dependent quality control pathways lead to convergent molecular and neurodevelopmental pathology. Elife. 2021;10.

18. Duncan JL, Pierce EA, Laster AM, Daiger SP, Birch DG, Ash JD, et al. Inherited Retinal Degenerations: Current Landscape and Knowledge Gaps. Transl Vis Sci Technol. 2018;7(4):6.

19. Zanni G, Kalscheuer VM, Friedrich A, Barresi S, Alfieri P, Di Capua M, et al. A Novel Mutation in RPL10 (Ribosomal Protein L10) Causes X-Linked Intellectual Disability, Cerebellar Hypoplasia, and Spondylo-Epiphyseal Dysplasia. Hum Mutat. 2015;36(12):1155–8.

20. Ishimura R, Nagy G, Dotu I, Zhou H, Yang XL, Schimmel P, et al. RNA function. Ribosome stalling induced by mutation of a CNS-specific tRNA causes neurodegeneration. Science. 2014;345(6195):455–9.

21. Terrey M, Adamson SI, Gibson AL, Deng T, Ishimura R, Chuang JH, et al. GTPBP1 resolves paused ribosomes to maintain neuronal homeostasis. Elife. 2020;9.

22. Yang L, Yan B, Qu L, Ren J, Li Q, Wang J, et al. IGF2BP3 Regulates TMA7-mediated Autophagy and Cisplatin Resistance in Laryngeal Cancer via m6A RNA Methylation. Int J Biol Sci. 2023;19(5):1382–400.

23. Juszkiewicz S, Slodkowicz G, Lin Z, Freire-Pritchett P, Peak-Chew SY, Hegde RS. Ribosome collisions trigger cis-acting feedback inhibition of translation initiation. Elife. 2020;9.

24. Sinha NK, Ordureau A, Best K, Saba JA, Zinshteyn B, Sundaramoorthy E, et al. EDF1 coordinates cellular responses to ribosome collisions. Elife. 2020;9.

25. Bryan JM, Fufa TD, Bharti K, Brooks BP, Hufnagel RB, McGaughey DM. Identifying core biological processes distinguishing human eye tissues with precise systems-level gene expression analyses and weighted correlation networks. Hum Mol Genet. 2018;27(19):3325–39.

26. Swamy V, McGaughey D. Eye in a Disk: eyeIntegration Human Pan-Eye and Body Transcriptome Database Version 1.0. Invest Ophthalmol Vis Sci. 2019;60(8):3236–46.

27. Evan, Basu A, Satija R, Nemesh J, Shekhar K, Goldman M, et al. Highly Parallel Genome-wide Expression Profiling of Individual Cells Using Nanoliter Droplets. Cell. 2015;161(5):1202–14.

28. Clark BS, Stein-O’Brien GL, Shiau F, Cannon GH, Davis-Marcisak E, Sherman T, et al. Single-Cell RNA-Seq Analysis of Retinal Development Identifies NFI Factors as Regulating Mitotic Exit and Late-Born Cell Specification. Neuron. 2019;102(6):1111–26.e5.

29. Hilal T, Yamamoto H, Loerke J, Burger J, Mielke T, Spahn CM. Structural insights into ribosomal rescue by Dom34 and Hbs1 at near-atomic resolution. Nat Commun. 2016;7:13521.

30. Doma MK, Parker R. Endonucleolytic cleavage of eukaryotic mRNAs with stalls in translation elongation. Nature. 2006;440(7083):561–4.

31. Tsuboi T, Kuroha K, Kudo K, Makino S, Inoue E, Kashima I, et al. Dom34:hbs1 plays a general role in quality-control systems by dissociation of a stalled ribosome at the 3’ end of aberrant mRNA. Mol Cell. 2012;46(4):518–29.

32. Izawa T, Tsuboi T, Kuroha K, Inada T, Nishikawa S, Endo T. Roles of dom34:hbs1 in nonstop protein clearance from translocators for normal organelle protein influx. Cell Rep. 2012;2(3):447–53.

33. Pisareva VP, Skabkin MA, Hellen CU, Pestova TV, Pisarev AV. Dissociation by Pelota, Hbs1 and ABCE1 of mammalian vacant 80S ribosomes and stalled elongation complexes. EMBO J. 2011;30(9):1804–17.

34. Jamar NH, Kritsiligkou P, Grant CM. The non-stop decay mRNA surveillance pathway is required for oxidative stress tolerance. Nucleic Acids Res. 2017;45(11):6881–93.

35. Guydosh NR, Kimmig P, Walter P, Green R. Regulated Ire1-dependent mRNA decay requires no-go mRNA degradation to maintain endoplasmic reticulum homeostasis in S. pombe. Elife. 2017;6.

36. Lewandowski D, Sander CL, Tworak A, Gao F, Xu Q, Skowronska-Krawczyk D. Dynamic lipid turnover in photoreceptors and retinal pigment epithelium throughout life. Prog Retin Eye Res. 2022;89:101037.

37. Chen HY, Kelley RA, Li T, Swaroop A. Primary cilia biogenesis and associated retinal ciliopathies. Semin Cell Dev Biol. 2021;110:70–88.

38. Barnes CL, Malhotra H, Calvert PD. Compartmentalization of Photoreceptor Sensory Cilia. Front Cell Dev Biol. 2021;9:636737.

39. Juszkiewicz S, Speldewinde SH, Wan L, Svejstrup JQ, Hegde RS. The ASC-1 Complex Disassembles Collided Ribosomes. Mol Cell. 2020;79(4):603–14 e8.

40. Hashimoto S, Sugiyama T, Yamazaki R, Nobuta R, Inada T. Identification of a novel trigger complex that facilitates ribosome-associated quality control in mammalian cells. Sci Rep. 2020;10(1):3422.

41. Juszkiewicz S, Chandrasekaran V, Lin Z, Kraatz S, Ramakrishnan V, Hegde RS. ZNF598 Is a Quality Control Sensor of Collided Ribosomes. Mol Cell. 2018;72(3):469–81 e7.

42. Ballabio E, Mariotti M, De Benedictis L, Maier JA. The dual role of endothelial differentiation-related factor-1 in the cytosol and nucleus: modulation by protein kinase A. Cell Mol Life Sci. 2004;61(9):1069–74.

43. Stoneley M, Harvey RF, Mulroney TE, Mordue R, Jukes-Jones R, Cain K, et al. Unresolved stalled ribosome complexes restrict cell-cycle progression after genotoxic stress. Mol Cell. 2022;82(8):1557–72 e7.

44. Dickinson ME, Flenniken AM, Ji X, Teboul L, Wong MD, White JK, et al. High-throughput discovery of novel developmental phenotypes. Nature. 2016;537(7621):508–14.

45. Mattapallil MJ, Wawrousek EF, Chan C-C, Zhao H, Roychoudhury J, Ferguson TA, et al. TheRd8Mutation of theCrb1Gene Is Present in Vendor Lines of C57BL/6N Mice and Embryonic Stem Cells, and Confounds Ocular Induced Mutant Phenotypes. Investigative Opthalmology & Visual Science. 2012;53(6):2921.

46. Livak KJ, Schmittgen TD. Analysis of relative gene expression data using real-time quantitative PCR and the 2(-Delta Delta C(T)) Method. Methods. 2001;25(4):402–8.

47. Wang Z, Liu CH, Huang S, Chen J. Assessment and Characterization of Hyaloid Vessels in Mice. J Vis Exp. 2019(147).

48. Gong Y, Li J, Sun Y, Fu Z, Liu C-H, Evans L, et al. Optimization of an Image-Guided Laser-Induced Choroidal Neovascularization Model in Mice. PLOS ONE. 2015;10(7):e0132643.

49. Navarrete-Perea J, Yu Q, Gygi SP, Paulo JA. Streamlined Tandem Mass Tag (SL-TMT) Protocol: An Efficient Strategy for Quantitative (Phospho)proteome Profiling Using Tandem Mass Tag-Synchronous Precursor Selection-MS3. Journal of Proteome Research. 2018;17(6):2226–36.

50. R Core Team. R: A language and environment for statistical computing. Vienna, Austria: R Foundation for Statistical Computing; 2022.

51. RStudio Team. RStudio: Integrated Development Environment for R. PBC, Boston, MA: RStudio; 2022.

52. C W. _cowplot: Streamlined Plot Theme and Plot Annotations for ‘ggplot2’_. R package 1.1.1; 2020.

53. Wickham H. ggplot2: Elegant Graphics for Data Analysis: Springer-Verlag New York; 2016.

54. Slowikowski K. _ggrepel: Automatically Position Non-Overlapping Text Labels with ‘ggplot2’_. R package version 0.9.2; 2022.

55. Warnes G BB, Bonebakker L, Gentleman R, Huber W, Liaw A, Lumley, T MM, Magnusson A, Moeller S, Schwartz M, Venables B. _gplots: Various R Programming Tools for Plotting Data_. R package version 3.1.3; 2022.

56. Galili T. dendextend: an R package for visualizing, adjusting and comparing trees of hierarchical clustering. Bioinformatics. 2015;31(22):3718–20.

57. Shannon P, Markiel A, Ozier O, Baliga NS, Wang JT, Ramage D, et al. Cytoscape: a software environment for integrated models of biomolecular interaction networks. Genome Res. 2003;13(11):2498–504.

58. Raudvere U, Kolberg L, Kuzmin I, Arak T, Adler P, Peterson H, et al. g:Profiler: a web server for functional enrichment analysis and conversions of gene lists (2019 update). Nucleic Acids Res. 2019;47(W1):W191–W8.

59. Subramanian A, Tamayo P, Mootha VK, Mukherjee S, Ebert BL, Gillette MA, et al. Gene set enrichment analysis: a knowledge-based approach for interpreting genome-wide expression profiles. Proc Natl Acad Sci U S A. 2005;102(43):15545–50.

60. Mootha VK, Lindgren CM, Eriksson KF, Subramanian A, Sihag S, Lehar J, et al. PGC-1alpha-responsive genes involved in oxidative phosphorylation are coordinately downregulated in human diabetes. Nat Genet. 2003;34(3):267–73.

61. Liberzon A, Birger C, Thorvaldsdottir H, Ghandi M, Mesirov JP, Tamayo P. The Molecular Signatures Database (MSigDB) hallmark gene set collection. Cell Syst. 2015;1(6):417–25.

62. Liberzon A, Subramanian A, Pinchback R, Thorvaldsdottir H, Tamayo P, Mesirov JP. Molecular signatures database (MSigDB) 3.0. Bioinformatics. 2011;27(12):1739–40.

63. Morris JH, Apeltsin L, Newman AM, Baumbach J, Wittkop T, Su G, et al. clusterMaker: a multi-algorithm clustering plugin for Cytoscape. BMC Bioinformatics. 2011;12:436.

64. Oesper L, Merico D, Isserlin R, Bader GD. WordCloud: a Cytoscape plugin to create a visual semantic summary of networks. Source Code Biol Med. 2011;6:7.

65. Merico D, Isserlin R, Stueker O, Emili A, Bader GD. Enrichment map: a network-based method for gene-set enrichment visualization and interpretation. PLoS One. 2010;5(11):e13984.

66. Reimand J, Isserlin R, Voisin V, Kucera M, Tannus-Lopes C, Rostamianfar A, et al. Pathway enrichment analysis and visualization of omics data using g:Profiler, GSEA, Cytoscape and EnrichmentMap. Nat Protoc. 2019;14(2):482–517.

67. Li Q, Lin J, Widrick JJ, Luo S, Li G, Zhang Y, et al. Dynamin-2 reduction rescues the skeletal myopathy of a SPEG-deficient mouse model. JCI Insight. 2022;7(15).

68. Perez-Riverol Y, Xu QW, Wang R, Uszkoreit J, Griss J, Sanchez A, et al. PRIDE Inspector Toolsuite: Moving Toward a Universal Visualization Tool for Proteomics Data Standard Formats and Quality Assessment of ProteomeXchange Datasets. Mol Cell Proteomics. 2016;15(1):305–17.

69. Deutsch EW, Bandeira N, Perez-Riverol Y, Sharma V, Carver JJ, Mendoza L, et al. The ProteomeXchange consortium at 10 years: 2023 update. Nucleic Acids Res. 2023;51(D1):D1539–D48.

70. Perez-Riverol Y, Bai J, Bandla C, Garcia-Seisdedos D, Hewapathirana S, Kamatchinathan S, et al. The PRIDE database resources in 2022: a hub for mass spectrometry-based proteomics evidences. Nucleic Acids Res. 2022;50(D1):D543–D52.

